# Amoeboid Cell Migration through Regular Arrays of Micropillars under Confinement

**DOI:** 10.1101/2022.04.08.487483

**Authors:** Zeinab Sadjadi, Doriane Vesperini, Annalena M. Laurent, Lena Barnefske, Emmanuel Terriac, Franziska Lautenschläger, Heiko Rieger

**Author notes:** Z. S. and D. V. contributed equally to this work. F. L. and H. R. contributed equally to this work.

## Abstract

Migrating cells often encounter a wide variety of topographic features—including the presence of obstacles—when navigating through crowded biological environments. Unravelling the impact of topography and crowding on the dynamics of cells is key to better understand many essential physiological processes such as the immune response. We study how migration and search efficiency of HL-60 cells differentiated into neutrophils in quasi two-dimensional environments are influenced by the lateral and vertical confinement and spatial arrangement of obstacles. A microfluidic device is designed to track the cells in confining geometries between two parallel plates with distance *h*, in which identical micropillars are arranged in regular pillar forests. We find that at each cell-pillar contact event, the cell spends a finite time near the pillar surface, which is independent of the height *h* and the interpillar spacing *e*. At low pillar density regime, the directional persistence of cells reduces with decreasing *h* or *e*, influencing their diffusivity and first-passage properties. The dynamics is strikingly different at high pillar density regime, where the cells are in simultaneous contact with more than one pillar; the cell velocity and persistence are distinctly higher compared to dilute pillar configurations with the same *h*. Our simulations reveal that the interplay between cell persistence and cell-pillar interactions can dramatically affect cell diffusivity and, thus, its first-passage properties.

## INTRODUCTION

Cell migration is essential for various physiological processes such as wound healing, morphogenesis, and immune responses [1–3]. Cells and other organisms can adapt their migration in response to different environmental cues such as gradients of chemical, electrical, or mechanical signals. Recently, the ability of migrating cells to sense and follow topographic environmental cues has attracted attention, the so-called *topotaxis* [4, 5]. Similar to other taxis phenomena, variations in topographic features of the surrounding environment— such as the spatial arrangement of obstacles, degree of lateral confinement, surface topography, etc.— can be exploited by biological organisms to navigate more efficiently [5– 10]. The idea of topotaxis can be utilized to conduct the migration of cells, e.g. by tuning spatial confinements or designing favorable arrangements of obstacles. To achieve an efficient topotaxis, however, a detailed understanding of the impact of crowding and confinement on different cell migration modes is required, which is currently lacking.

Amoeboid migration is a fast cell migration mode, which relies on friction instead of adhesion [6, 11, 12]. Various cell types— including immune, stem, and metastatic tumor cells— exhibit this mode of migration. Amoeboid migration of immune cells has been of particular interest [13–16], since one of their main functions is to explore the environment to detect pathogens. To reach this aim, they need to pass through highly confined tissues and extracellular matrices, whereby the degree on confinement depends on the type of tissue and organ they are migrating in. Cell migration experiments through arrays of micropillars *in vitro* [5–7] have revealed that two regimes of pillar densities can be distinguished: high density regime— where the cell is often attached to several pillars simultaneously and experiences a directed pillar-to-pillar type of motion— and low density regime— where the cell usually contacts only one or two pillars simultaneously—. The cells are relatively faster at high density regime, but the velocity is not considerably affected by pillar density within this regime [5, 6]. It has been shown that mesenchymal cells at low adhesion benefit from lateral confinement by switching to an amoeboid-like migration and move faster [17]; however, it is unclear how the degree of the imposed lateral confinement influences the migratory behavior in the amoeboid mode of migration in quasi-2D environments.

Additionally, the role of cell-obstacle interactions on cell dynamics, navigation, and search efficiency at low obstacle densities has not been well understood yet. The nature of the interactions with obstacles is expected to crucially affect the dynamics of the moving objects. It can occasionally lead to an increased diffusivity or search efficiency. For instance, the presence of bystander cells accelerates the migration of natural killer cells [18]. Also, bacteria can increase their diffusivity by sliding along the surface of micropillars [19–21]. Nevertheless, obstacles often slow down the dynamics of moving objects due to steric, hydrodynamic, or frictional effects. Microalgae scatter of surfaces by pushing against them with their flagella [22–24]. Scattering from obstacles— e.g. through classical specular reflection as in the Lorentz gas model— randomizes the trajectory; the asymptotic diffusion constant decreases monotonically with increasing the volume fraction occupied by the obstacles [25–33]. Moreover, moving objects such as swimming bacteria can be hydrodynamically captured near the surface of sufficiently large micropillars [21]. The critical trapping distance *δ*_*c*_— below which the swimmer is captured near the pillar surface— and the trapping time *τ*_*c*_ increase with the size of micropillars [20]. Consistently, the contact times of same-sized bystander cells or polystyrene beads with migrating natural killer cells have been comparable [18]. However, less is known about the trapping by obstacles for amoeboid migration in confined geometries, where the cells form frictional contacts with surfaces.

To understand how the combined effects of scattering from and trapping by obstacles govern the dynamics of migrating cells, random-walk models can be a powerful tool to untangle complex cell migratory behavior from the experimental data. Stochastic two-state models consisting of altering phases of fast and slow motions, such as run-and-tumble or run-and-pause dynamics, have been widely employed to describe locomotive patterns in biological systems [34–39]. A proper numerical model, however, needs to be capable of capturing the topographical features of the problem.

In this work, we study the topographical influence of the environment on *in vitro* amoeboid cell migration in regular arrays of micropillars. The cells move in quasi-2D environments confined between two parallel plates. By varying the plate-plate distance (vertical confinement), the interpillar spacing (lateral confinement), and the spatial arrangement of pillars, we track the trajectories of the cells to demonstrate their dynamics and to determine the characteristics of their interactions with pillars— including the thickness *δ* of the cell-pillar contact zone and the contact time *τ*_*c*_—. The influence of such cell-pillar interactions on search efficiency of the cells is nontrivial: although spending time in the vicinity of pillars slows the spreading and enhances the search time, the extent of the contact zone around the pillars reduces the effective search area which has an opposite effect. Using numerical simulations, we clarify how the interplay between the vertical confinement, spatial pillar distribution, and cell-pillar interaction governs the dynamics and first-passage properties of the migrating cells.

## MATERIALS AND METHODS

### Cells

HL-60 acute promyelocytic cell line was cultured in Roswell Park Memorial Institute medium (RPMI-1640, Gibco) supplemented with 10% fetal bovin serum (Fisher Scientific), 1% Glutamax (Fisher Scientific) and 1% penicillin/streptomycin (Gibco). HL-60 cells were differentiated into neutrophils with 1.3% DMSO for 3 days before performing the experiments.

### Pillar forest geometries

The pillar forest chambers were designed with Autodesk Inventor [40]. The device consisted of a cell loading inlet and a tracking area (see Fig. 1 for details). The tracking area consisted of six vertically stacked chambers, each with a dimension of 500 × 500 *µ*m^2^. The plate-plate distance was determined by the pillar height *h*. The chambers were filled with pillars with a diameter *d* and an inter-pillar spacing distance *e*, organized either in a triangular (T) or square (S) lattice. The resulting chambers had different pillar densities, named as *dense, intermediate* and *sparse*. Three sets of devices with increasing *h* were named as D1, D2, and D3, respectively. For the triangular lattice we had an extra dense device D4 with *e* = 5 *µ*m, named as *packed* device. The chambers alternated from square to triangular lattice with a decreasing density. The geometrical information of the pillar forests is summarized in Table I. See also a full list of the parameters used in the paper in the Suppl. Table S1.

**FIG. 1.**
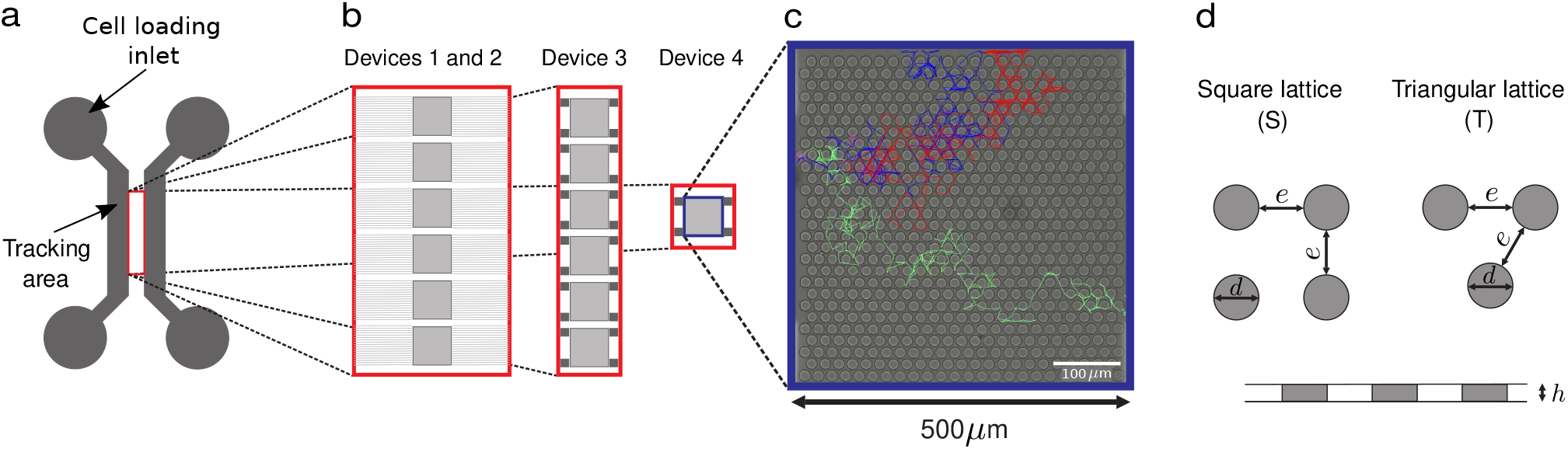
Sketch of the experimental devices. (a) General view of the cell loading channels. (b) Zoom of the tracking area for each device. (c) Example of the bright field image of device D4 with 3 cell tracks of different colors. (d) Definition of the pillar diameter *d*, pillar-pillar spacing *e*, and plate-plate distance *h*, for square (S) and triangular (T) lattices (see the summary of the geometrical parameters in Table I).

**TABLE 1.** Geometrical parameters of the experimental devices. The plate-plate distance *h*, pillar diameter *d*, and interpillar spacing *e* are given in *µ*m for triangular (T) or square (S) lattices.

| Device |  | D1 |  |  |  | D2 |  |  |  | D3 |  |  |  | D4 |  |  |  |
| --- | --- | --- | --- | --- | --- | --- | --- | --- | --- | --- | --- | --- | --- | --- | --- | --- | --- |
| Method |  | Photolithography |  |  |  |  |  |  |  | Two-photon lithography |  |  |  |  |  |  |  |
| Geometry | | $d$ | $e$ | $h$ | $\lambda$ | $d$ | $e$ | $h$ | $\lambda$ | $d$ | $e$ | $h$ | $\lambda$ | $d$ | $e$ | $h$ | $\lambda$ |
| Sparse | T1 | 11 | 49 | 3.5 | 0.18 | 13 | 47 | 5 | 0.22 | 15 | 45 | 6 | 0.25 | - | - | - | - |
|  | S1 |  |  |  |  |  |  |  |  |  |  |  |  |  |  |  |  |
| Intermediate | T2 | 11 | 29 | 3.5 | 0.27 | 13 | 28 | 5 | 0.31 | 15 | 25 | 6 | 0.37 | - | - | - | - |
|  | S2 |  |  |  |  |  |  |  |  |  |  |  |  |  |  |  |  |
| Dense | T3 | 12 | 18 | 3.5 | 0.40 | 13 | 17 | 5 | 0.43 | 15 | 15 | 6 | 0.50 | - | - | - | - |
|  | S3 |  |  |  |  |  |  |  |  |  |  |  |  |  |  |  |  |
| Packed | T4 | - | - | - | - | - | - | - | - | - | - | - | - | 15 | 5 | 4 | 0.75 |

### Production of the wafers

The tracking area of the devices D1 and D2 consisted of six chambers, connected to the cell loading channel of 900 *µ*m width and 50 *µ*m height via twenty small channels of 10 *µ*m width and 3.5 *µ*m or 5 *µ*m height (see Fig. 1). They were fabricated using the standard photolithography technique, processing guidelines from Microchem, in two steps. Briefly, a 4-inch silicon wafer was covered with a first layer (tracking area) of SU8-3005, spin coated at 500 rpm for 15 s followed by 4000 rpm for 40 s or 3000 rpm for 30 s, for getting desirable heights (respectively 3.5 *µ*m and 5 *µ*m). Then the wafer was soft baked for 2 min at 95^◦^C, and exposed to UV light (UV-KUB-2, Kloe, France) through a mask with an illumination of 50% for 7 s and 8 s, respectively. The wafer was then post baked for 2 min at 95^◦^C, developed in a developer solution for 1 min and rinsed with isopropanol. The second layer (cell loading inlets and channels) of the master fabrication was performed using SU8-3025, spin coated at 500 rpm for 15 s and 1500 rpm for 45 s. Then the wafer was soft baked for 2 min at 95^◦^C, and exposed to UV light through a mask with an illumination of 50% during 32 s. The wafer was then post baked for 5 min at 95^◦^C, developed in a developer solution for 8 min and rinsed with isopropanol. The tracking area of the devices D3 and D4 consisted of six chambers and one chamber, respectively, which were connected to the cell loading channel of 900 *µ*m width and 100 *µ*m height via square channels of 100 × 100 × 100 *µ*m^3^ placed at each corner of each chamber (see Fig. 1). They were printed by two-photon lithography with Nanoscribe GT+ (Nanoscribe, Germany) with IP-S resin (Nanoscribe, Germany) on ITO-coated glass substrates using a 25× objective. A laser intensity of 150 mW and a writing speed of 100 mm/s was applied.

First washed with PGMEA, exchanged against isoporpanol, post cured for 5 minutes under 200 W UV radiation (OmniCure Series 1500, IGB-Tech GmbH). The samples were carefully dried under nitrogen stream. In order to reduce printing time the chambers were fully printed of high accuracy and the cell loading inlet was printed with shell and scaffold printing mode where a posting curing process was necessary.

Production of the microfabricated devices Microfabricated devices were replica molded into silicone rubber (RTV615, Momentive Performance Materials, USA) using soft lithography. Briefly, the silicon rubber was cast onto the wafer, degassed and polymerized at 75^◦^C for 2 hours. The resulting devices were peeled off and sealed in 35 mm glass bottom cell culture dishes (World Precision Instruments, Sarasota, FL, USA) using plasma surface activation.

### Experimental setup

Prior to experiment, the assembled structure was coated with 100 *µ*g.mL^−1^ poly-L-lysine (20 kDa) grafted with polyethylene glycol (2 kDa) (PLL-PEG) (Sigma-Aldrich, St Louis, MO, USA) for 30 min at room temperature. Cell nuclei were stained with 200 ng.mL^−1^ Hoechst 34580 (Sigma Aldrich, St Louis, USA), for 30 min before being platted at the cell loading channel with a concentration of 5 × 10^3^ cells.mL^−1^. When cells started to fill the chamber, the cell culture dish was filled with RPMI medium and kept at 37^◦^C for at least 30 min before starting the experiment. Fluorescent images of cell nuclei and bright field images of the pillar chamber were recorded using a EMCCD camera (Andor Technology, UK) with a physical pixel size of 0.65 *µ*m and a binning 2 × 2, mounted on a Nikon Eclipse Ti epifluorescent microscope, at a 10 × magnification and 0.5 NA over 12 h with a frame rate of 2 min. The cells were kept at constant atmosphere of 37^◦^C and 5% CO_2_ (Okolab, Italy) during the entire experiment. To minimize bleaching effect, the exposure times were kept at 100 ms for the fluorescent images and 20 ms for the bright field images.

### Data analysis

Cell trajectories were analyzed using ImageJ plugin TrackMate. We excluded the trajectories of dying and dividing cells. The maximum tracking time was 700 min, however, we excluded the first 100 min of all tracks until the cells reached the bulk of the chambers. Since the cells entered the camera field at different times, we shifted the starting time of all trajectories to have all cells starting at the same time, which is *t* = 100 min in real time in our experiments. Each trajectory consisted of a set of (*x, y*) positions, recorded after successive time intervals Δ*t* = 2 min. Every two successive recorded positions were used to calculate the instantaneous velocity and every three of them to extract the local turning angle *ϕ*. A small *ϕ* corresponds to a highly persistent motion, i.e. moving nearly along the previous direction of motion. On the other hand, *ϕ* approaches *π* when the direction of motion is nearly reversed. We quantified the local cell persistence with cos *ϕ*, ranging from − 1 for forward motion to 1 for backward turning. The mean local persistence was then obtained as ℛ = cos *ϕ* [41], with ℛ ∈ [− 1, 1]. All statistical quantities were calculated for each geometry, by adding up all trajectories in the corresponding experiments. Minimum number of cell trajectories analyzed in each experiment as well as the number of experiments performed for each geometry are summarized in Suppl. Table S2. The mean square displacement (MSD) was calculated as MSD(*t*) = ⟨ *r*(*t*)^2^ ⟨ − ⟨ *r*(*t*) ⟩ ^2^⟩, with ⟨ … ⟩ being the average over all cell trajectories in one chamber.

### Simulation method

Monte Carlo simulations were performed to study the migration of cells through a two-dimensional medium consisting of circular obstacles. To mimic each experiment, the corresponding experimental distributions of velocity and persistence, the setup dimensions, and the positions and distances between the pillars served as input for simulations. An ensemble of 10^4^ persistent random walkers started their motion from a random position on the left border (as in the experiments) and with a random shooting angle into the simulation box with periodic boundary conditions. We also performed control simulations in a simulation box of 200 × 200 *µ*m consisting of a lattice of *N × N* circular obstacles (*N∈* [4, 5, 6, 7, 8]) with diameter *d* = 10 *µ*m. We used an ensemble of 10^6^ persistent random walkers, which started their motion from a random position with a random direction. Each walker spent a waiting time *τ*_*c*_ when reaching a distance *δ* from the surface of an obstacle. We systematically varied the persistence of the random walker and *τ*_*c*_ and *δ* values, beyond the available range of these parameters in our experiments.

## RESULTS

In order to understand the influence of vertical confinement and pillar density on amoeboid cell migration, we use three devices D1, D2 and D3 with the plate-plate distance *h* = 3.5, 5 and 6 *µ*m, respectively. Each of these devices contains pillars of diameter *d* arranged on square (S) or triangular (T) lattice configurations with different pillar spacing *e* (see the “Materials and methods” section, Table I, and Fig. 1 for details of geometrical properties). Because of using different techniques to produce our devices, *d* and *e* parameters— which determine the fraction of the occupied space by pillars— vary from chamber to chamber. Thus, we characterize the pillar density with the relative pillar size 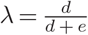 to take the effects of both parameters *d* and *e* into account. *λ* is a dimensionless quantity ranging from 0 for point-like pillars (*d* = 0) to 1 for the maximum possible pillar size (*e* = 0), i.e. when the neighboring pillars are in contact with each other. The mean diameter of differentiated HL-60 cells in our experiments is around 10 *µ*m, which is smaller than the interpillar distance in all chambers of devices D1, D2, and D3. However, we also construct a highly dense device D4 with *e* = 5 *µ*m and *h* = 4 *µ*m; thus, here the cells can be vertically and laterally confined from several sides.

### Cell dynamics

Cells enter the chambers from one side and move through pillars. The mean instantaneous velocity *v* of cells in devices D1, D2 and D3 does not systematically depend on the pillar density, lattice type, or plate-plate distance, as shown in Fig. 2a. However, the cells are significantly faster in device D4, with a mean velocity *v* = 5.01 ± 0.02 *µ*m*/*min. The velocity distribution *P* (*v*)— presented in Fig. 2b for triangular and in Suppl. Fig. S1a for square lattices— similarly shows no trend in terms of *h* or *λ*. The tail of *P* (*v*) decays faster than exponential for all chambers, with a relatively slower decay for device D4.

**FIG. 2.**
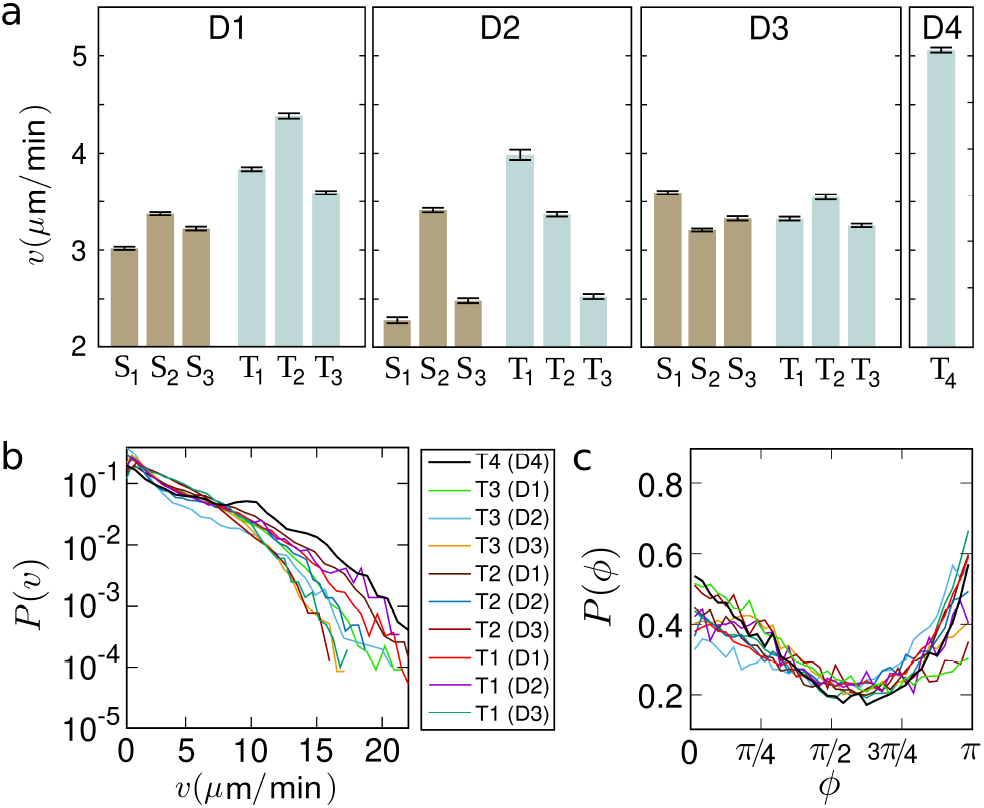
(a) Mean cell velocity *v* in different devices D. The results are separately shown for different interpillar distances in triangular (T) or square (S) lattice configurations. The error bars indicate the standard errors of the means over all instantaneous velocities in each chamber. The total number of cell trajectories and experiments per chamber are given in Suppl. Table S2. (b) Velocity distribution *P* (*v*) in log-lin scale for all triangular configurations. The characteristics of each configuration are given in Table I. (c) Turning-angle distribution *P* (*ϕ*) for all triangular configurations. All line colors are as in panel (b).

The turning-angle distribution *P* (*ϕ*) of the cells is shown in Fig. 2c for all triangular lattices (see the square lattice data in Suppl. Fig. S1b). In all cases, *P* (*ϕ*) develops two peaks around *ϕ* = 0 and *π*, reflecting that the motion in near forward or backward directions are more probable. We quantify the mean local persistence of cells in each chamber by the dimensionless parameter ℛ = ⟨ cos *ϕ* ⟩ [41], where ⟨ … ⟩ denotes averaging over all cell trajectories in one chamber. ℛ ranges from 1 for pure localization to 0 for diffusion and 1 for ballistic motion (see the “Materials and methods” section for data analysis details).

Figure 3 shows the mean local persistence ℛ versus the relative pillar size *λ* for various devices. ℛ reduces with decreasing *h* in both lattice types. Also, at the given vertical confinements *h* = 5 *µ*m or 6 *µ*m (devices D2 and D3), ℛ reduces with increasing the density of pillars. However, this trend is not observed in device D1 with *h* = 3.5 *µ*m, where the cells move nearly diffusively, i.e. with no effective propulsion, independent of the pillar density. Cell persistence in device D4 is distinctly high, despite having a considerably higher pillar density.

**FIG. 3.**
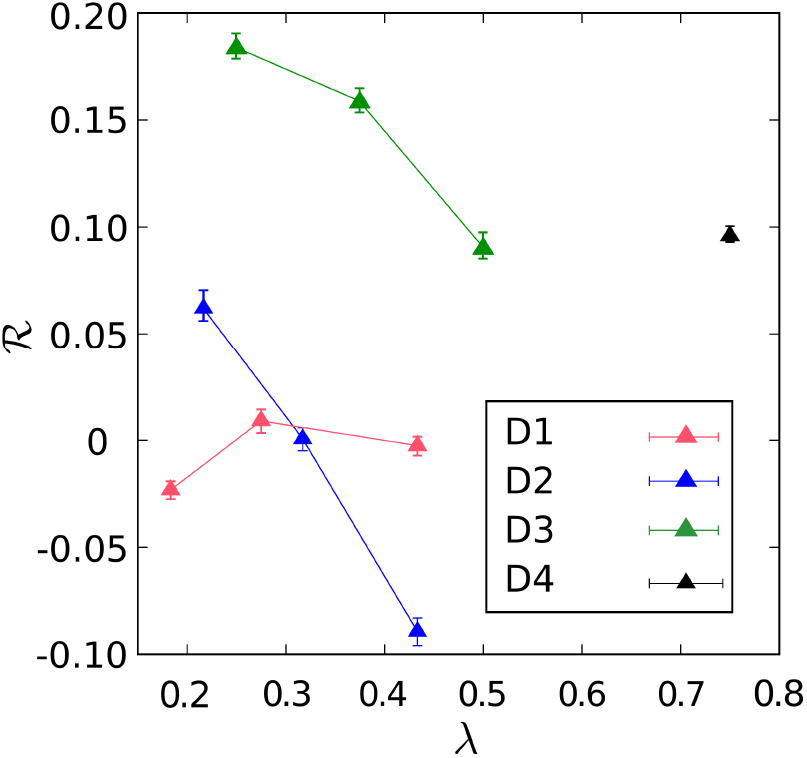
Mean local persistence ℛ of the cells versus the relative pillar size *λ* = *d /*(*d* + *e*) for different chambers with triangular lattices of micropillars. The error bars indicate the standard errors of the means. For the square lattice data see Suppl. Fig. S2.

In addition to the local persistence parameter ℛ, we also quantify the cell persistence with a parameter *γ*(*t*), which reflects the overall curvature of cell trajectories until time *t*. We obtain the path length *ℓ*(*t*) and the net displacement of the cell *ℓ*_net_(*t*) at time *t* with respect to the starting time. By averaging 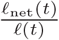 over all cell trajectories, the overall persistence of the trajectories until time *t* can be quantified as 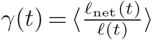. As shown in Fig. 4a, *γ*(*t*) behaves similarly to ℛ versus *λ* or *h*; it increases with decreasing *λ* (from T3 to T1) or increasing *h* (from D1 to D3). The time evolution of *γ*(*t*) in an ordinary diffusion with constant velocity follows *γ*(*t*) ∼ *t*^−1*/*2^, since the path length grows linearly with time while the net displacement is proportional to the square root of the MSD, i.e. 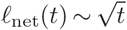. For comparison, in a persistent random walk with the same constant velocity, *γ*(*t*) is initially larger than in the ordinary diffusion, but it similarly decays as *t*^−1*/*2^ at long times.

**FIG. 4.**
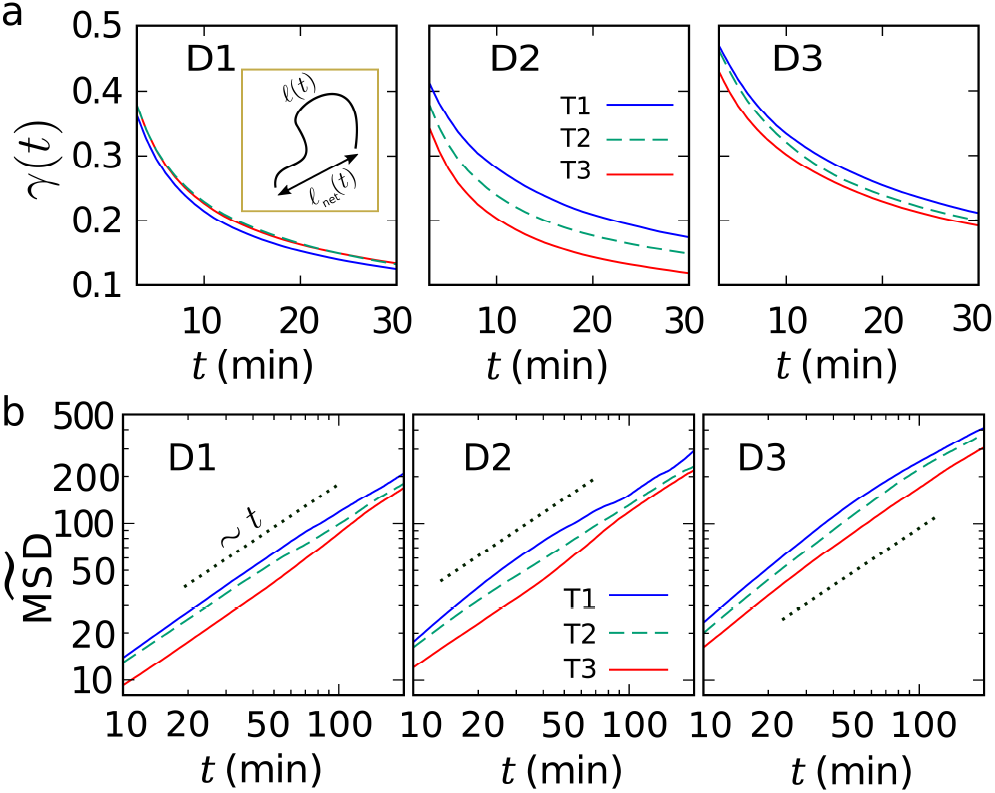
Time evolution of (a) the overall persistence *γ*(*t*) of cell trajectories and (b) the scaled MSD of cells, for triangular lattices with different chamber thickness and pillar density. The inset of panel (a) depicts the path length *ℓ*(*t*) and the net displacement *ℓ*_net_(*t*) of a cell trajectory. The dotted lines in (b) represent normal diffusion and serve as a guide to the eye. For the square lattice data see Suppl. Fig. S3.

We compare the MSD of the cells in different devices in Fig. 4b. The main goal in each panel of this figure is to understand the role of pillar density on the diffusivity. Nevertheless, the mean and variance of velocity vary from experiment to experiment, thus, a direct comparison of the MSD curves is not informative. It is known that the MSD of a persistently moving object in a uniform space depends on the velocity moments as [42]

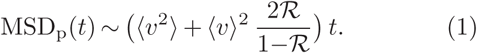

To be able to compare the MSD from different experiments, we rescale them by MSD_p_(*t*) from Eq. (1) using the corresponding experimental values. The resulting scaled 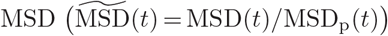, shown in Fig. 4b, reveals that increasing the pillar density (i.e. from T1 to T3) leads to a lower diffusivity in all devices. The relative decay range of the diffusion constant from T1 to T3 is around 19%, 32%, and 27% for devices D1, D2, and D3, respectively.

### Escape times

Next, we address how the geometrical characteristics of the pillar forest influence the first-passage properties of the cells. We estimate the mean first-passage time as the mean residence time *t*_esc_ which is spent by the cell in the area confined between adjacent pillars (see the grey zone in Fig. 5a and Ref. [[5]]). Indeed, *t*_esc_ is the time which takes for a cell to escape a local trap formed by adjacent pillars and move to the next trap. The area of each trap zone is 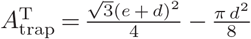 or 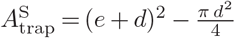 for square or triangular lattices, respectively. With increasing *λ, A*_trap_ gets smaller, thus, the escape time *t*_esc_ expectedly reduces, as shown in Fig. 5c. At the same *λ, t*_esc_ is smaller for triangular lattices compared to square configurations since 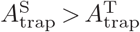. To correct for this effect, we divide *t*_esc_ by *A*_trap_ in Fig. 5d. The resulting escape time per unit area increases with *λ*, which shows that the increase of obstacles per unit area strengthens the trapping effect and enhances the escape time.

**FIG. 5.**
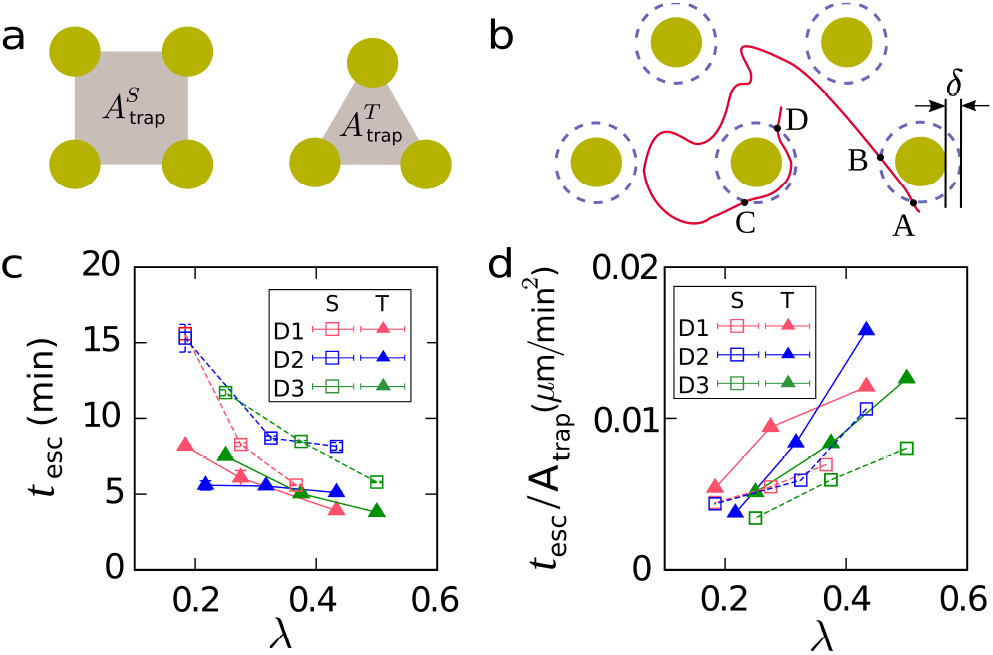
(a) Schematic drawing of the trap zone (grey region), i.e. the region confined between adjacent pillars. The corresponding areas are denoted with 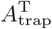 and 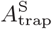 in triangular and square lattices, respectively. (b) Schematic example of a cell trajectory which visits the contact zones twice. The corresponding contact times *τ*_c_ are *t*_*A*→*B*_ and *t*_*C*→*D*_, and it spends *τ*_b_ = *tB*→*C* in the bulk. The dashed circles indicate the borders of the contact zones, with the contact distance *δ* from the pillar surfaces. (c) Mean escape time *t*_esc_ versus the relative pillar size *λ* for different chambers. (d) Mean escape time, scaled by the trap zone area *A*_trap_, versus *λ* for different chambers.

### Cell-pillar interactions

To characterize the cell-pillar interaction, we measure the time spent by cells in the vicinity of pillars. We define a contact zone around each pillar as the region within a distance *δ* from the pillar surface (see Fig. 5b), and define a contact event when a cell surface enters this zone. We measure the contact time *τ*_*c*_ as the time spent by a cell in a contact zone in each contact event (a contact event occurs when the distance between the cell nucleus and the center position of the pillar falls below the sum of the cell radius, *δ*, and the pillar radius). For different choices of *δ*, we measure *τ*_*c*_ for all cell trajectories belonging to each chamber. The typical result is presented in Fig. 6a for device D3 with *e* = 45 *µ*m. It can be seen that below a critical distance *δ*_*c*_ ≈ 4 *µ*m, the contact time *τ*_*c*_ is independent of the choice of *δ*, evidencing the formation of the cell-pillar contact. We choose a contact distance *δ* = 2 *µ*m within the plateau regime (i.e. *δ <δ*_*c*_) for all chambers, and measure the resulting contact time *τ*_*c*_ in different experiments. Except for two experiments (D2,T1 and D3,T3; see Table I), the resulting mean contact time *τ*_*c*_ is around 3.8 0.2 min for all chambers, independent of *h, λ*, or lattice type (Fig. 6b). The longer contact time in two of the experiments originates from the extremely long stay of some cells in the vicinity of pillars. We speculate that these rare events are caused by abnormal or dying cells or local defects on pillar surfaces.

**FIG. 6.**
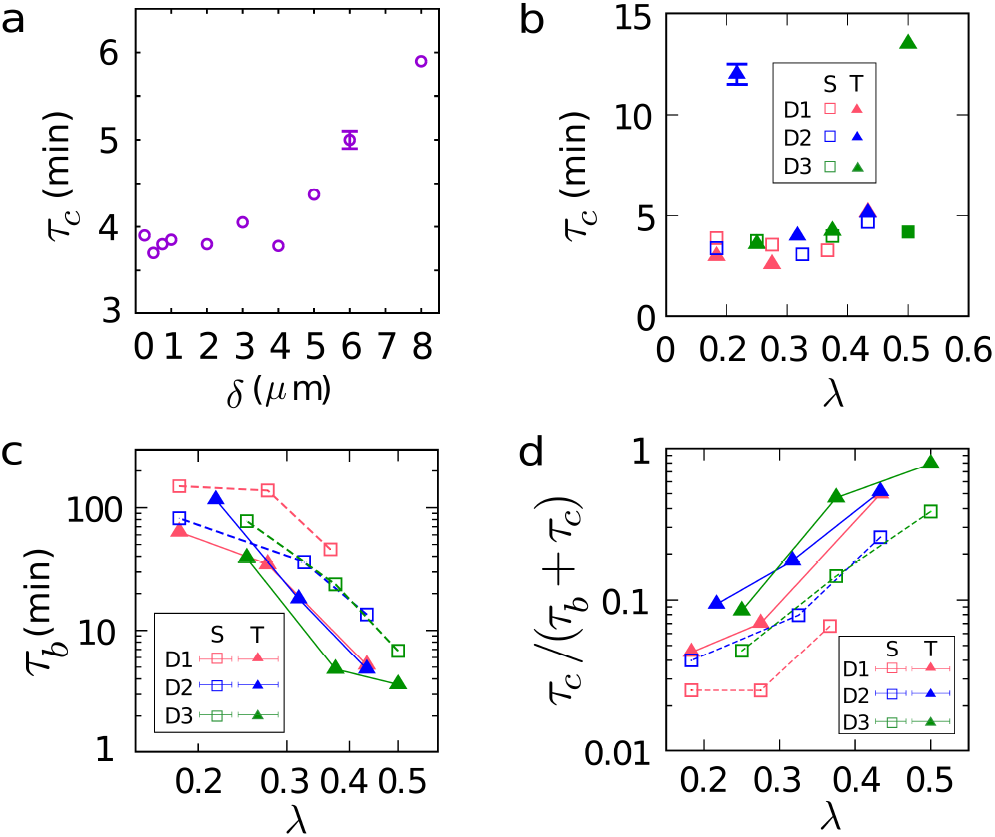
(a) Contact time *τ*_c_ versus the thickness *δ* of the contact zone, for device D3 with *e* = 45 *µ*m. (b-d) Mean contact time *τ*_c_ (b), mean bulk time *τ*_b_ (c), and the fraction of time spent in the vicinity of pillars 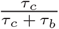 (d) versus the relative pillar size *λ*, for different chambers. The single error bars shown in panels (a,b) represent the typical estimated errors for all data. Panels (c) and (d) are presented in log-log and log-lin scales, respectively.

We similarly introduce a bulk time *τ*_*b*_ as the duration of time that a cell spends in the bulk of the pillar forest between two successive contact events. The mean value of the bulk time *τ*_*b*_ is presented in Fig. 6c for different chambers. *τ*_*b*_ decreases with increasing *λ*, since the available bulk area decreases and the cells visit the pillars more frequently. Also, the larger values of *τ*_*b*_ in square lattices compared to triangular ones is due to the larger available area in the square configuration compared to the triangular lattice with the same *λ* (i.e. the same interpillar spacing *e*; see Fig. 5a). In Fig. 6d, the fraction of time spent in the vicinity of pillars is shown. This fraction increases with the density of pillars, as the relative contribution of the contact events increases. It can be also seen that the relative contact time is the least in the highly vertically confined device D1.

### Numerical results

According to our experimental observations, the directional persistence ℛ of the cells in amoeboid migration through the micropillar arrays is relatively weak and depends on the vertical confinement *h* in a given geometry of pillars. Motivated by this, we model the migration of cells with a persistent random walk (PRW) in a two-dimensional medium, containing circular obstacles. Thus, the impact of the vertical confinement *h* is considered by the persistence of the persistent ℛ random walker in our model. We first validate our numerical model by comparing it to the experimental data. For each experiment, the corresponding geometrical quantities *d, e*, and pillar positions are used as input for simulations. Persistent random walkers with velocity and turning-angle distributions compatible with each experiment (as presented in Figs. 2b, 2c and Suppl. Fig. S1) are considered. The random walker halts for a mean contact time *τ*_*c*_ = 3.8 min in the vicinity of pillars, when it enters a contact zone with *δ* = 2 *µ*m, compatible with the results of the previous subsection. The details of the simulation method are presented in the “Materials and methods” section. Moreover, an ordinary random walk (ORW) model with and without a constant residence time *τ*_*c*_ near the obstacles is considered for comparison. The constant velocity of the walker in the ORW model is chosen to be the mean velocity of the cells in each experiment and the turning-angle distribution is uniform.

Our goal is to understand the navigation and search abilities of migrating cells in the amoeboid mode. The mean-first-passage time is conversely related to the diffusion constant *D* of the searcher [43–45]; thus, we focus on the asymptotic regime of cell dynamics and compare the numerically obtained MSD with the experimental results. As typical examples with different persistence ℛ, we present in Fig. 7 the results for experiments D3,S2 and D2,S2 (see Table I), which have a finite or nearly zero persistence, respectively. The results of the PRW model with empirical input matches very well with the experimental results in both cases. The ORW model with constant velocity neglects the cell persistence and the variance of the velocity distribution, which leads to a smaller MSD compared to the PRW model [42]. Taking the waiting time at contact events into account further lowers the ORW model curve in Fig. 7. We checked that the PRW model satisfactory captures the time evolution of the MSD in other chambers as well.

**FIG. 7.**
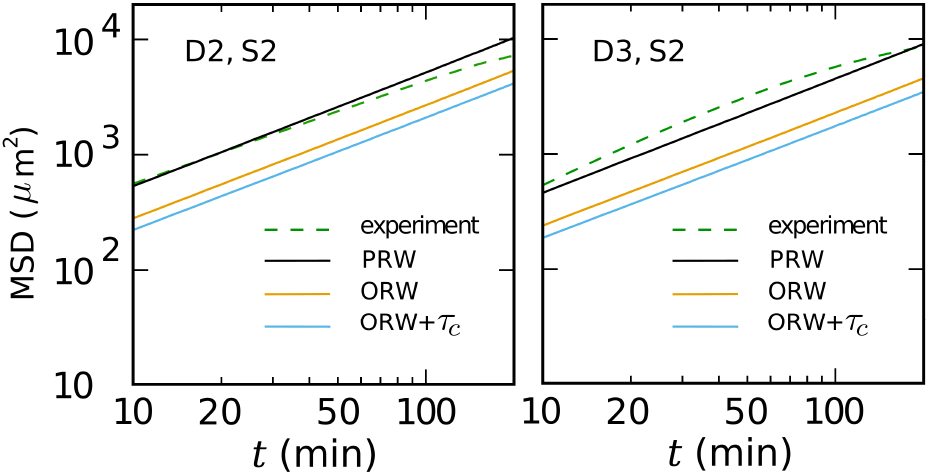
(a) Time evolution of the mean square displacement in the asymptotic regime. A comparison is made between the simulation and experimental results in experiment D2,S2 (left) or D3,T2 (right); see Table I for the geometrical parameters of each case. The simulation results are presented for a PRW model with experimental data served as input for simulations, and an ordinary random walk (ORW) model with a constant velocity, with or without a constant residence time near the pillars.

In order to gain more insight into the role of the key parameters on the amoeboid cell migration, we perform extensive Monte Carlo simulations of the PRW model. The walker moves through a square lattice of circular obstacles and stays in the contact zones around the obstacles for a finite time. See the “Materials and methods” section for details. In our control simulations, a reference set of parameters— *λ* = 0.25, ℛ = 0 (pure diffusion), *τ*_*c*_ = 2 min, and *δ* = 2 *µ*m— is chosen. The velocity in all simulations has a uniform distribution with the mean velocity of 3 *µ*m*/*min. We systematically vary each parameter beyond the available range in our experiments, while other parameters are kept fixed at their reference values. We particularly focus here on how cell persistence, crowding by obstacles, contact-zone size, and contact time influence the asymptotic diffusion constant *D* of the cell.

The dependence of *D* on the relative pillar size *λ* is shown in Fig. 8a. *D* expectedly decreases with increasing pillar density; within the experimental variation range of *λ*, the diffusion constant decays by approximately 30% in simulations, which is comparable with the experimental observations.

**FIG. 8.**
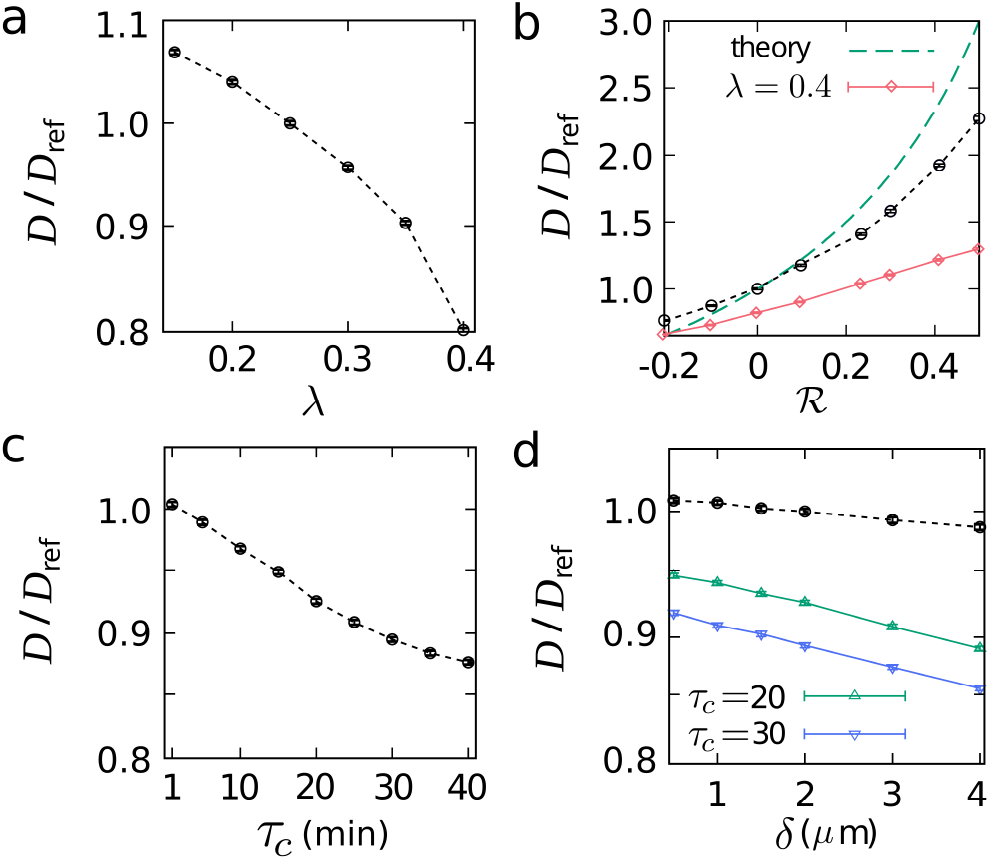
Simulation results for diffusion constant *D* of persistent random walkers in a 2D obstacle forest. The diffusion constant for the reference set of parameters (see text) is denoted by *D*_ref_. The diffusion constant *D*, scaled by *D*_ref_, is shown in terms of (a) the relative pillar size *λ*, (b) mean persistence R, (c) cell-pillar contact time *τ*_c_, and (d) cell-pillar contact zone size *δ*. Except for the varied parameter in each panel, the rest of the parameters are kept fixed at their reference values, unless specified otherwise. The dashed line in panel (b) represents the theoretical prediction Eq. (2).

Next we vary the persistence ℛ of the random walker. For this purpose, the width of a uniform turning-angle distribution *P* (*ϕ*) around the forward direction of motion is varied in simulations. Increasing ℛ results in a larger diffusion constant *D* (Fig. 8b). For comparison, *D* of a persistent random walker in the absence of pillars grows with ℛ as [42]

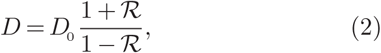

with *D*0 being the diffusion constant of a normal diffusion with the same mean velocity as the persistent random walker. The presence of obstacles reduces *D* and limits its variation range. We also present the results for a denser system with *λ* = 0.4 in Fig. 8b; the variation range of *D* is even further limited. The impact of obstacles on *D* is more pronounced at larger values of ℛ. In the small persistence regime (as for the migrating cells in our experiments), the influence of obstacles on *D* gets relatively weaker. Interestingly, for subdiffusive dynamics (i.e. ℛ *<* 0), the presence of obstacles can even increase *D*, as the obstacles randomize the path of the slowly spreading anti-persistent random walker.

Finally, we investigate the role of the characteristics of cell-pillar contacts, including the contact time *τ*_*c*_ and contact-zone size *δ* in Figs. 8c,d. Within the experimental range of *τ*_*c*_ (see Fig. 6b), variation range of *D* is small. However, larger values of *τ*_*c*_ enhance the contribution of the waiting events and visibly affect *D*. By varying the contact-zone size *δ* at the reference contact time *τ*_*c*_ = 2 min, the walker contacts the obstacles more frequently. The influence on *D* is however weak, as shown in Fig. 8d. The reason is that the reference contact time is so small that the contribution of the increased number of contact events is negligible. By increasing *τ*_*c*_ to 20 min or 30 min, variation of *δ* leads to stronger changes in *D*.

## DISCUSSION

We have studied the *in vitro* amoeboid migration of HL-60 cells differentiated into neutrophils in quasi-2D confined geometries containing regularly arranged cylindrical micropillars. The spacing between identical pillars and their lattice type of arrangement have been varied to study their impact on the amoeboid migration in the dilute regime of obstacles. In this regime, the pillars act as scatterers and randomize the cell trajectory. Moreover, the cells get locally trapped between adjacent pillars, which slows the cell dynamics. In our experiments, the interpillar distance is chosen to be larger than the typical size of HL-60 cells, which is around 10 *µ*m. There is, however, one exceptional chamber (device D4) with a high pillar density in such a way that the interpillar spacing is in the range of the size of the cell nucleus. The cell is then in contact with several pillars simultaneously. In this case, the cell benefits from a directed pillar-to-pillar type of motion to increase its persistence and velocity (Figs. 2, 3). Nevertheless, we mainly focus on the dilute pillar density regime. Here, decreasing the interpillar distance reduces the persistence and asymptotic diffusion constant *D* of migrating cells (Figs. 3, 4). It is known that increasing the obstacle density in regular arrangements of symmetric obstacles or random configurations of them slows the particle dynamics due to increasing effects of reorientations by obstacles and/or trapping in local cages between them[25–33, 46]. Our numerical simulations reveal that with doubling the relative pillar size *λ, D* can decrease even more than 30% (Fig. 8a). As *D* is conversely related to the mean-first-passage time [43– 45], the search time of the cell— e.g. the mean escape time from the local cages formed by adjacent pillars— is expected to increase with *λ*. This is confirmed by our experimental results; doubling *λ* increases the escape time per unit area by a factor of about 2 (Fig. 5d). We note that the asymptotic diffusion limit can be inaccessible in experiments, e.g., due to high concentration of obstacles [27]. While the searcher may not explore the entire space in such cases, it can still explore the local environment through an anomalous diffusive dynamics [29].

So far, it has been unclear how vertical confinement influences amoeboid cell migration. Although contact with surfaces is required to initiate and maintain the amoeboid migration, our results reveal that being too squeezed between parallel plates impairs the migration in low pillar density regime. The cell size in our experiments has been larger than the chamber height *h* in all devices. Upon decreasing the plate-plate distances below *h* 4 ≈ *µ*m, the cells practically lose their migration ability and just diffuse (Fig. 3). It remains for future studies how the generation of the biomechanical forces required for amoeboid migration depends on the vertical confinement of the cell. The simulation results in Fig. 8b reveal that the variation of cell persistence ℛ (upon changing the chamber height *h*) can significantly affect the asymptotic diffusion constant. The impact of ℛ (equivalently *h*) on the cell dynamics however weakens with increasing the obstacle density in the environment.

By investigating the cell-pillar interactions in our experiments, we have determined the mean contact-zone size *δ* and contact time *τ*_*c*_ of cells with pillars. The question remains how far these contact events influence the dynamics and first-passage properties of migrating cells. To gain more insight into the role of *τ*_*c*_ and *δ* on cell migration, we have varied these parameters in numerical simulations. Importantly, we find that the contact time of our cells with pillars is too short to be able to affect the cell dynamics and the asymptotic diffusion constant *D*. Because of the short waiting times at contact events, extending the contact zone area by increasing *δ* has a negligible influence on *D* (black dashed curve in Fig. 8d). However, by increasing *τ*_*c*_ beyond the plateau regime at small times, the decay of *D* accelerates; additionally, increasing the contact-zone size *δ* at a longer reference *τ*_*c*_ leads to considerable changes in *D*.

To conclude, we find that the obstacles act as scatterers to randomize the dynamics of migrating cells, when we focus on the dilute regime. We also find that, the adjacent obstacles create local traps which further slow the cell dynamics. Therefore, in this regime, obstacles impair the amoeboid cell migration rather than facilitate it. In contrast, simultaneous contacts with several obstacles in the dense regime help the cells to move forward from obstacle to obstacle, in agreement with [7]. Our results highlight the importance of the vertical confinement on amoeboid cell migration in the dilute regime; the cells lose their migration ability when extremely squeezed between parallel plates. In future, our work can be used to understand the origin of different migration behavior of immune cells in different tissues and organs. We are also convinced that our data will help to mimic and control amoeboid migration in vitro by choosing appropriate confinement parameters.

## SUPPLEMENTARY INFORMATION

Supplementary information accompanies this paper, including three figures.

## AUTHOR CONTRIBUTIONS

F.L. and H.R. designed the research. D.V., A.M.L., L.B., E.T. and F.L. performed experiments. D.V. and Z.S. analyzed data. Z.S. performed numerical simulations. Z.S. wrote the manuscript. All authors revised the manuscript.

## ACKNOWLEDGEMENTS

We thank Galia Montalvo for reading the manuscript. We acknowledge support from the Deutsche Forschungsgemeinschaft (DFG) through the collaborative research center SFB 1027.

