## Supplementary material for "Amoeboid Cell Migration through Regular Arrays of Micropillars under Confinement": Figures and Tables

### Supplementary Information to Amoeboid Cell Migration through Regular Arrays of Micropillars under Lateral Confinement

**Table S1:** Summary of all parameters used in trecordedhe paper.

| | |
|---|---|
| $d$ | Pillar diameter |
| $e$ | Interpillar spacing |
| $\lambda$ | Relative pillar size $\lambda = d/(d+e)$ |
| $h$ | Height of pillars, i.e. plate-plate distance of a device |
| $A_{\text{trap}}$ | Area of each trap zone, depicted in Fig. 5a |
| $\phi$ | Turning angle of a cell in two successive recorded frames |
| $P(\phi)$ | Probability distribution of turning angles |
| $\mathscr{R}$ | Mean local persistence $\mathscr{R} = \langle cos\phi \rangle$, averaged over all turning angles of trajectories in a geometry |
| $v$ | Mean instantaneous velocity of all trajectories recorded in a geometry |
| $P(v)$ | Probability distribution of $v$ |
| $t_{esc}$ | Mean residence time which is spent by the cell in the trap zone $A_{\text{trap}}$ |
| $\delta$ | A parameter which defines the contact zone around each pillar |
| $\tau_c$ | Mean contact time spent by the cell in contact with a pillar in a single contact |
| $\tau_b$ | Mean bulk time spent by the cell in the bulk between two of its successive contacts with pillars |
| $\ell(t)$ | Path length traveled by the cell until time $t$ |
| $\ell_{net}(t)$ | Net displacement of the cell until time $t$ |
| $\gamma(t)$ | Averaged $\frac{\ell_{\text{net}}(t)}{\ell(t)}$ over all cell trajectories in each geometry |

**Table S2:** Number of experiments $N_{\text{exp}}$, and the minimum number of trajectories $N_{\text{traj}}$ analyzed per geometry.

| Device | | D1 | | D2 | | D3 | | D4 | |
|---|---|---|---|---|---|---|---|---|---|
| | | $N_{\text{exp}}$ | $N_{\text{traj}}$ | $N_{\text{exp}}$ | $N_{\text{traj}}$ | $N_{\text{exp}}$ | $N_{\text{traj}}$ | $N_{\text{exp}}$ | $N_{\text{traj}}$ |
| Sparse | T1 | 10 | 100 | 5 | 20 | 4 | 50 | - | - |
| | S1 | 7 | 100 | 5 | 20 | 3 | 60 | - | - |
| Intermediate | T2 | 9 | 60 | 6 | 80 | 5 | 20 | - | - |
| | S2 | 7 | 100 | 4 | 60 | 3 | 40 | - | - |
| Dense | T3 | 11 | 100 | 4 | 60 | 6 | 40 | - | - |
| | S3 | 7 | 100 | 4 | 60 | 3 | 40 | - | - |
| Packed | T4 | - | - | - | - | - | - | 46 | 180 |

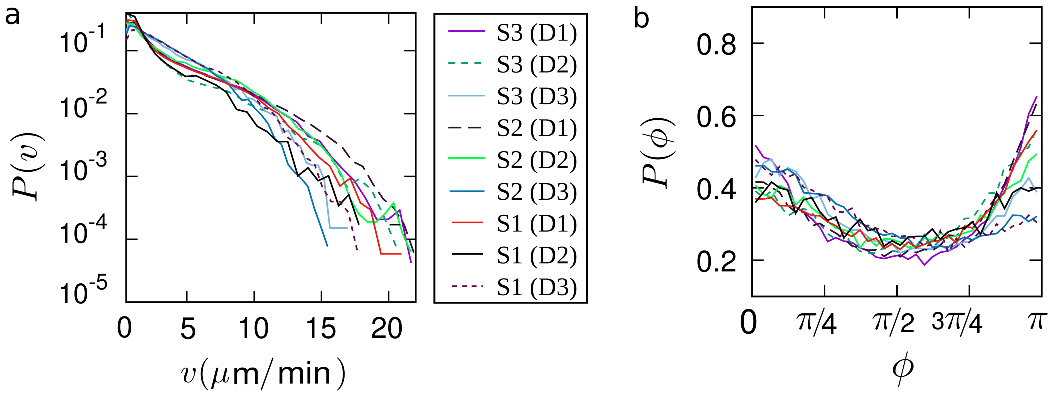


**Figure S1:** (a) Velocity distribution $P(v)$ in log-lin scale for all square configurations. The characteristics of each configuration are given in Table I. (b) Turning-angle distribution $P(\phi)$ for all square configurations. All line types and colors are as in panel (a).

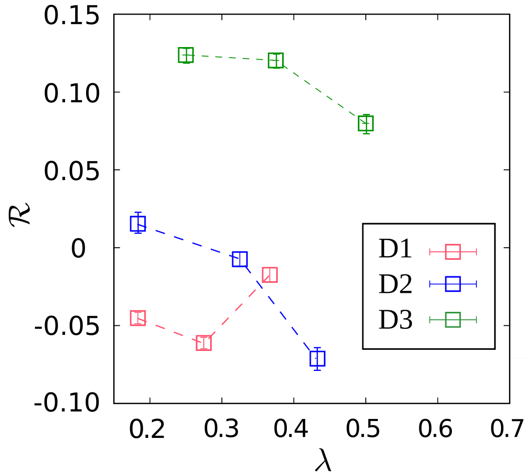


**Figure S2:** Mean local persistence $\mathscr{R}$ of the cells versus the relative pillar size $\lambda = d/(d+e)$ for different chambers with square lattices of micropillars. The error bars indicate the standard errors of the means.

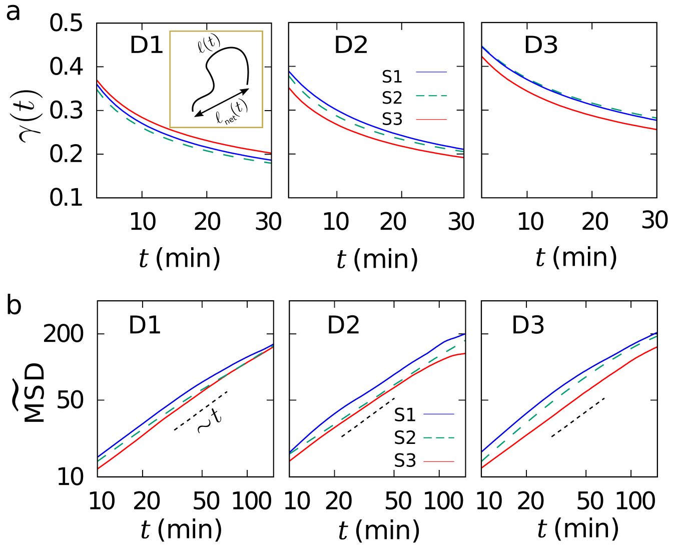


**Figure S3:** Time evolution of (a) the overall persistence $\gamma(t)$ of cell trajectories and (b) the scaled MSD of cells, for square lattices with different chamber thickness and pillar density. The inset of panel (a) depicts the path length $\ell(t)$ and the net displacement $\ell_{\mathrm{net}}(t)$ of a cell trajectory. The dotted lines in (b) represent normal diffusion and serve as a guide to the eye.
